# Assessing neuromodulation effects of theta burst stimulation to the prefrontal cortex using TMS-evoked potentials

**DOI:** 10.1101/2021.06.21.449219

**Authors:** Adriano H. Moffa, Stevan Nikolin, Donel Martin, Colleen Loo, Tjeerd W. Boonstra

## Abstract

Theta burst stimulation (TBS), a form of repetitive transcranial magnetic stimulation (TMS), is capable of non-invasively modulating cortical excitability. TBS is gaining popularity as a therapeutic tool for psychiatric disorders such as depression, in which the dorsolateral prefrontal cortex (DLPFC) is the main therapeutic target. However, the neuromodulatory effects of TBS on prefrontal regions remain unclear. An emerging tool to assess neuromodulation in non-motor regions is concurrent transcranial magnetic stimulation and electroencephalography (TMS-EEG) to measure TMS-evoked potentials (TEPs). We assessed twenty-four healthy participants (13 males, mean age 25.2±9.9 years) following intermittent TBS, continuous TBS, and sham applied to the left DLPFC using a double-blinded crossover design. TEPs were obtained at baseline and 2-, 15-, and 30-min post-stimulation. Four TEP components (N40, P60, N100 and P200) were analysed using mixed effects repeated measures models (MRMM). Results indicate no significant effects for any assessed components (all p>.05). The largest effect size (Cohen’s d = −0.5) comparing iTBS and sham was obtained for the N100 component at 15 minutes post-stimulation. This result was in the same direction but smaller than found in previous studies, suggesting that the true effect size may be lower than previously reported. Accurate estimates of the effects sizes and inter-individual heterogeneity will critically inform clinical applications using TEPs to assess the neuromodulatory effects of TBS.

## 1. Introduction

Transcranial magnetic stimulation (TMS) is a form of non-invasive brain stimulation (NIBS) that uses focal electromagnetic fields to depolarise cortical neurons at depths of 2-3 cm (Wagner et al., 2009). A variation of repetitive TMS (rTMS) called theta-burst stimulation (TBS) involves the delivery of 50Hz triplet bursts of magnetic pulses repeated at a rate of 5Hz corresponding to the theta rhythm (Huang et al., 2005). Intermittent TBS (iTBS) has been shown to increase the excitability of the stimulated motor neurons (Huang et al., 2005, Di Lazzaro et al., 2008). Conversely, continuous TBS (cTBS) can be delivered to decrease cortical excitability (Di Lazzaro et al., 2005, Wischnewski and Schutter, 2015). Meta-analytical data suggest that the effects of TBS on motor cortical excitability may be equivalent to, if not greater than, those produced with standard rTMS protocols (Chung et al., 2016). TBS may also produce longer aftereffects, even with the use of relatively lower intensities (Nyffeler et al., 2006, Yang et al., 2015). Notwithstanding, there is limited knowledge about the neurophysiological mechanisms of TBS in brain regions outside of the motor cortex, e.g., the prefrontal cortex, a common target for TBS clinical interventions in neuropsychiatric conditions such as depression (Machado et al., 2013, Desmyter et al., 2016, Blumberger et al., 2018). Understanding the neurophysiological effects of prefrontal TBS stimulation may allow further refinement of the technique for therapeutic purposes.

Recent technological advances in hardware and data processing algorithms have enabled the possibility of assessing NIBS neuromodulatory effects at non-motor cortical sites using a combination of TMS and simultaneous electroencephalography (EEG) (Daskalakis et al., 2012). The method involves the delivery of single-pulse TMS to evoke a series of time-locked positive and negative deflections in the EEG traces (e.g. N40, P60, N100, P200), known as TMS-evoked EEG potentials (TEPs) (Ilmoniemi et al., 1997). Of these, the N100 deflection is considered the most robust (Noda et al., 2016) and reliable TEP component (Lioumis et al., 2009, Kerwin et al., 2018), with high sensitivity to detect small changes in cortical excitability (Nikulin et al., 2003). This makes the N100 component the best candidate for assessing the neuromodulatory effects of NIBS techniques, including TBS. Previous studies have assessed TEPs to explore the effects of TBS on the motor cortex and the cerebellum with mixed findings (see Tremblay et al. (2019) for a review). In the first experiment, which applied TBS to the left dorsolateral prefrontal cortex (DLPFC) of 10 healthy subjects, iTBS was shown to significantly increase N100 amplitudes in left frontal electrodes compared to sham, but no difference to cTBS nor between cTBS and sham was found (Chung et al., 2017). The same group also showed an inverse U-shaped relationship between stimulation intensity and modulatory aftereffects, with iTBS applied at 75% of resting motor threshold (RMT) yielding the most significant changes from pre-iTBS compared to both 100% and 50% RMT (Chung et al., 2018b). Compared to sham iTBS administered to age and gender-matched participants, an increased N100 amplitude was only observed with 75% iTBS in a fronto-central cluster. However, a more recent non-controlled study using a similar experimental design and larger sample size (n=20) did not show significant differences in N100 amplitude change from baseline between iTBS delivered at 30 Hz, 50 Hz and the subjects’ individualised theta-gamma coupling frequency (Chung et al., 2019). Moreover, inconsistent with previous findings, a decreased N100 amplitude was observed after individualised iTBS.

Given these discrepancies in results from previous studies (Tremblay et al., 2019) and considerable concerns in the literature regarding the validity of TEPs as a biomarker of neural function (Conde et al., 2019), we performed a conceptual replication of the first TBS-TEP study where TBS was administered to the left DLPFC to assess the neuromodulation effects of iTBS and cTBS using TEPs. Based on the study of Chung et al. (2017), we expected that iTBS would have a cortical excitatory effect, reflected in increased N100 amplitudes post stimulation compared to sham. By obtaining further evidence that TEPs can detect the neuromodulatory effects of prefrontal TBS stimulation, we aimed to evaluate the clinical utility of this technique.

## 2. Methods

### 2.1. Participants

Twenty-four healthy participants (13 males, mean age 25.2±9.9 years, range from 18 to 65) were included in the study, of which 21 were naïve to TMS. All were right-handed, as confirmed by the Edinburgh Handedness Inventory (Oldfield, 1971). The experimental procedure was approved by the Ethics committee of the University of New South Wales (HC17765). According to the declaration of Helsinki, all participants provided written informed consent prior to the start of the experiment. In addition, participants reported no history of neurological illness or brain injury, were non-smokers and not using any medication acting on the central nervous system at the time of the study.

### 2.2. Sample size calculation

The first study exploring the effects of TBS on the DLPFC found a significant difference between iTBS and sham in the relative change of N100 amplitude from pre-to post-TBS (Chung et al., 2017). Using the From_R2D2 effect size conversion algorithm (Lakens, 2013), we calculated the effect size presented in Chung et al. (2017) for the comparison of N100 amplitudes between sham and iTBS. This indicated an effect size of η^2^=0.36 and r=0.6, which corresponds to a large effect of Cohen’s d=1.5. Using a conservative approach, we chose to power our study to detect a moderately sized effect (Cohen’s d = 0.6) (Cohen, 1988) for the difference between iTBS and sham conditions in a predefined region of interest (ROI) over the left prefrontal cortex. Using the G*Power software (Faul et al., 2009), we calculated a sample size of 24 (power of 80% and an alpha value of 0.05).

### 2.3. Experimental design

This study is the first part of a larger study examining the effects of TBS on cortical excitability in a group of healthy participants. In this first part, each subject participated in three sessions (iTBS, cTBS and sham) in a double-blinded crossover design. The order of sessions was randomised across participants, and sessions were held at approximately the same time of the day with at least a one-week gap to avoid carryover effects. In each session, cortical excitability was assessed in blocks of 100 single TMS pulses recorded pre-TBS and at 2-, 15-, and 30-min post-TBS (Pre, T2, T15 and T30, respectively). In addition, eyes-open resting-state EEG was recorded for 4 minutes at the start of the session and after all four blocks of single-pulse TMS (Figure 1). During resting-state recordings, participants were instructed to keep their eyes open, look at a fixation cross presented in the middle of the screen and stay relaxed. The analysis of resting-state data will be reported elsewhere.

**Fig. 1.**
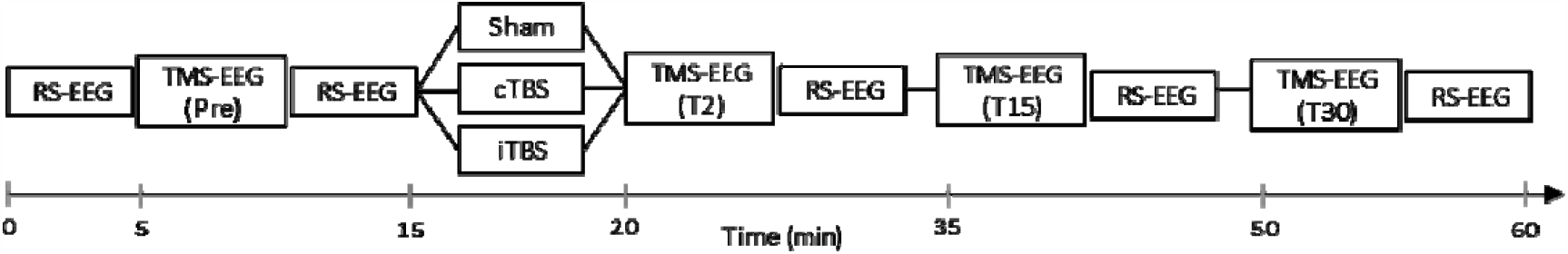
Experimental design/study timeline. First, open-eye resting-state electroencephalography (RS-EEG) was performed, followed by 100 single TMS pulses and the recording of the TEPs (TMS-EEG, Pre) and second block of RS-EEG. During the stimulation block, intermittent theta-burst stimulation (iTBS), continuous theta-burst stimulation (cTBS) or sham was administered in a randomized order in three separate sessions. TMS-EEG was repeated at T2 (2min), T15 (15 min) and T30 (30 min) after the stimulation block, each of them followed by RS-EEG.

### 2.4. Transcranial magnetic stimulation

Single-pulse stimulation and TBS were administered with a MagPro X100 (MagVenture A/S, Farum, Denmark) stimulator using an MC-B70 figure-of-8 coil oriented to the parasagittal plane using biphasic pulses. The centre of the TMS coil was positioned over the F3 electrode position using a 5-mm customised 3D printed spacer placed between the coil and the scalp to avoid contact with electrodes, minimising post-pulse artifacts, electrode movement and bone-conducted auditory input. The spacer base was marked on the cap and used as a reference for coil repositioning. A similar procedure was shown to improve consistency in coil location and angle within- and between-sessions when neuronavigation was not available (Rogasch et al., 2013, Chung et al., 2019).

Resting motor threshold (RMT) was determined using the Conforto et al. (2004) procedure by applying TMS over the cap with the spacer (see Supplementary Material S1).

To generate TEPs, participants received 100 single pulses (4s±10% jitter inter-pulse interval) per block delivered at a stimulation intensity of 120% RMT.

#### 2.4.1. Theta burst stimulation

Consistent with the original TBS protocol proposed by Huang et al. (2005), bursts of 3 stimuli at 50 Hz were repeated at intervals of 200 ms (i.e., 5Hz). The iTBS protocol involved a 2 s train of TBS repeated every 10 s for a total of 190 s (600 pulses), and cTBS consisted of a 40 s train of uninterrupted TBS (600 pulses). The intensity used for both active TBS conditions was 75% of RMT, as this intensity yielded the most pronounced neurophysiological changes for iTBS (Chung et al., 2018b). Sham iTBS was applied in half of the participants and sham cTBS for the other half. Sham TBS consisted of an inactive coil positioned on the head (at the same position as the active conditions) and a second active coil positioned 20 cm from the back of the head, facing away from it, with an increased stimulation output of 20% to compensate for the attenuation of the sound due to the additional distance from the ear. This sham condition was designed to avoid any direct cortical stimulation while matching the auditory input due to the TMS clicking sound. To minimise auditory evoked potentials from the clicking sound, participants listened to white noise masking played through in-ear earphones, individually adjusted until they could not hear the TMS clicking sound (single-pulse TMS at 120% RMT) or reached their upper threshold for comfort. In addition, earmuffs were placed over the earphones to further attenuate TMS pulse volume during the TMS-EEG blocks.

### 2.5. Electroencephalography (EEG)

#### 2.5.1. EEG recording

Continuous EEG data were acquired using a TMSi Refa amplifier (TMS International, Oldenzaal, Netherlands) and a 64-channel cap (EASYCAP Gmbh, Herrsching, Germany) with sintered, interrupted disk, Ag-AgCl TMS-compatible electrodes. To monitor eye blinks, electrooculography (EOG) was recorded using four additional electrodes, two positioned above and below the right eye, and the other two positioned directly lateral to the outer canthi of both eyes. Electrocardiography (ECG) was also recorded. Electrodes were grounded to FPz, and impedance levels were kept at <50 kΩ throughout the experiment, which is well below 1% of the input impedance (100MΩ) of the EEG amplifier (Metting van Rijn et al., 1991). EEG data were sampled at 2048 Hz. The trigger pulses controlling the TMS device were recorded together with the EEG to enable event-related averaging.

### 2.6. TMS-EEG data analysis 2.6.1 EEG pre-processing

EEG data were cleaned and analysed offline blind to the experimental conditions using a combination of open-source toolboxes: Fieldtrip (Oostenveld et al., 2011), EEGLAB (Delorme and Makeig, 2004), TESA (v0.1.0) (Rogasch et al., 2017) and custom scripts on the MATLAB platform (R2017b, The MathWorks, USA).

The cleaning parameters and procedures were based on the methods described in (Rogasch et al., 2017). First, data were epoched relative to the TMS pulse (−1000 to 1000 ms). Then, electrodes where the TMS artifact exceeded the maximum absolute value of the range of the amplifier (10^7 μV) were removed and linearly interpolated from neighbouring electrodes. On average, 2.0±1.7 channels were rejected in the cTBS condition, 2.1±1.8 channels in the iTBS condition and 2.0±1.8 channels in the sham condition. Trials in which the kurtosis exceeded 5 SD from the mean were excluded. The remaining trials were visually inspected to eliminate trials with excessive noise (bursts of muscle activity, electrode artifacts). On average, 11.2±12.5 trials were rejected in the cTBS condition, 12.5±6.1 trials in the iTBS condition and 11.5±6.6 trials in the sham condition per block of 100 trials.

EEG traces were detrended, and baseline corrected relative to pre-TMS data (−500 to −50 ms). Line noise (50 Hz) was removed using linear regression by fitting and subtracting a sine wave from the EEG time courses. Data between −5 and 10 ms around the TMS pulse were removed, and the first round of independent component analysis (ICA) was performed using the TESA *compselect* function to eliminate components containing eye blinks. On average, 5.0±3.3 components were removed at this stage. Next, TMS-muscle and decay artifacts were removed using a method proposed by Freche et al. (2018), in which a physical model of skin impedance is used to remove the artifactual signal without the degradation of the neural signal. This is achieved by first fitting analytic expressions (a power law in time rather than the commonly assumed exponential capacitive discharge curve) to the most negative and positive EEG signal deflections to obtain regression fit parameters and then removing the artifact from the data by subtraction (Freche et al., 2018).

Following removal of the TMS-muscle and decay artifacts, EEG data from before and after the TMS stimulus were filtered separately using a bandpass filter (1–90Hz). A more lateral stimulation site such as F3 can induce muscle activation and electrode movements (Huang and Mouraux, 2015, Ilmoniemi and Kičić, 2010), so a second round of ICA was performed to remove this as well as components associated with blinks, eye movement, persistent muscle activity, decay artifacts and electrode noise. With all four experimental blocks concatenated, a total of 14.0±3.1 components were removed at this stage. Removed channels were interpolated, and data were re-referenced to the common average reference. Finally, trials were separately averaged within each block to obtain the TEPs.

#### 2.6.2 TMS-evoked potentials (TEP)

Changes in cortical reactivity were assessed in a ROI over the left DLPFC to assess the local effects of TBS. We used an average of channels F1, F3, FC1 and FC3 as our ROI, similar to Chung et al. (2017). TEPs commonly show four separate peaks: N40, P60, N100 and P200 (Fig. 3). The latencies of the TEP peaks were identified on an individual basis within predefined time windows as the minimum amplitude between 30–50 ms (N40) and 90–140 ms (N100), and the maximum amplitude between 55–75 ms (P60) and 160–240 ms (P200) on the mean of the four ROI electrodes. The amplitude of each peak was then calculated as the average signal between ±5 ms of the previously identified peak latency (Chung et al., 2017).

**Fig. 2.**
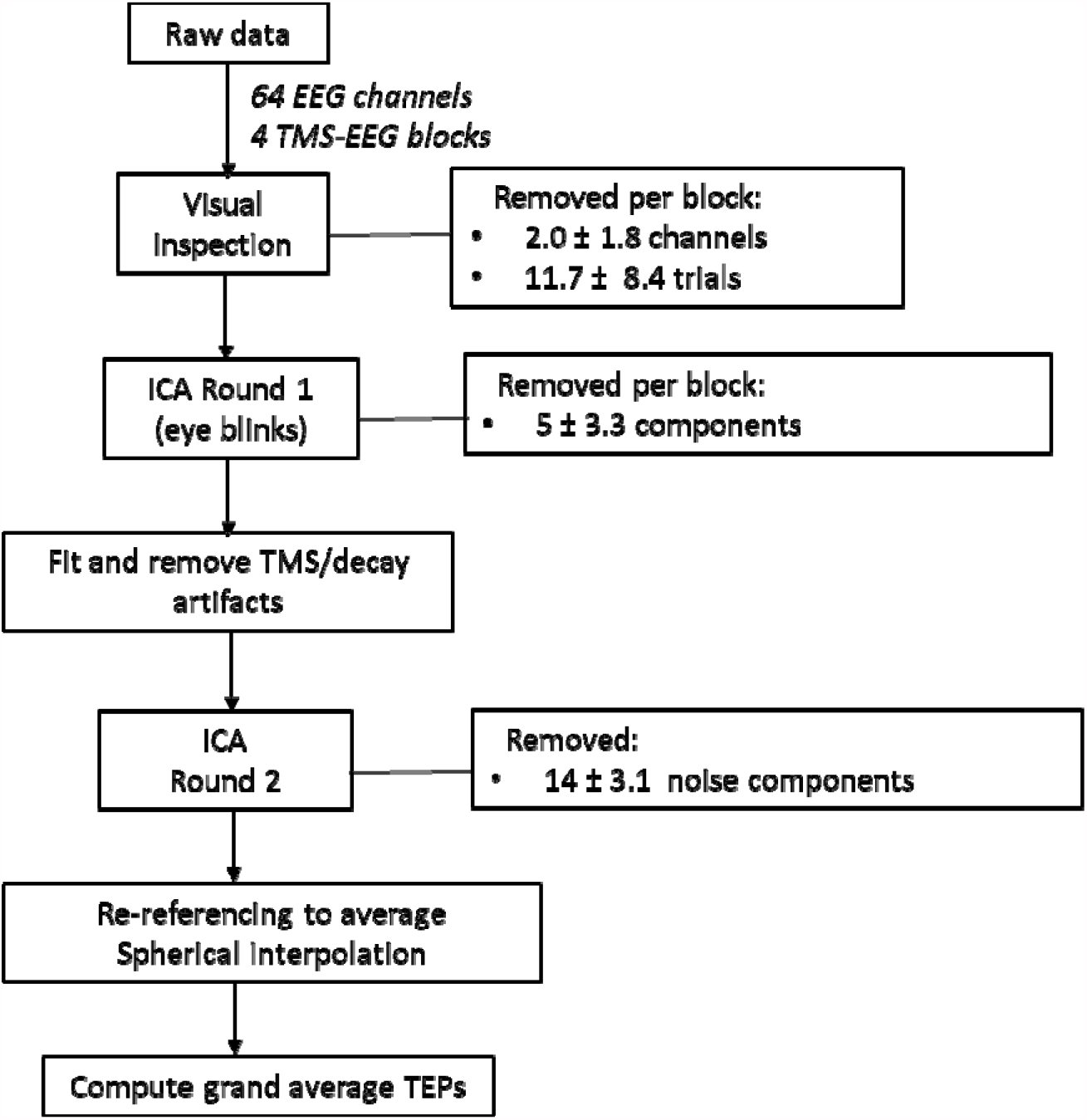
Data processing flowchart. TMS-EEG: transcranial magnetic stimulation-electroencephalography; ICA: independent component analysis; TMS: transcranial magnetic stimulation; TEPs: TMS-evoked potentials.

**Fig. 3.**
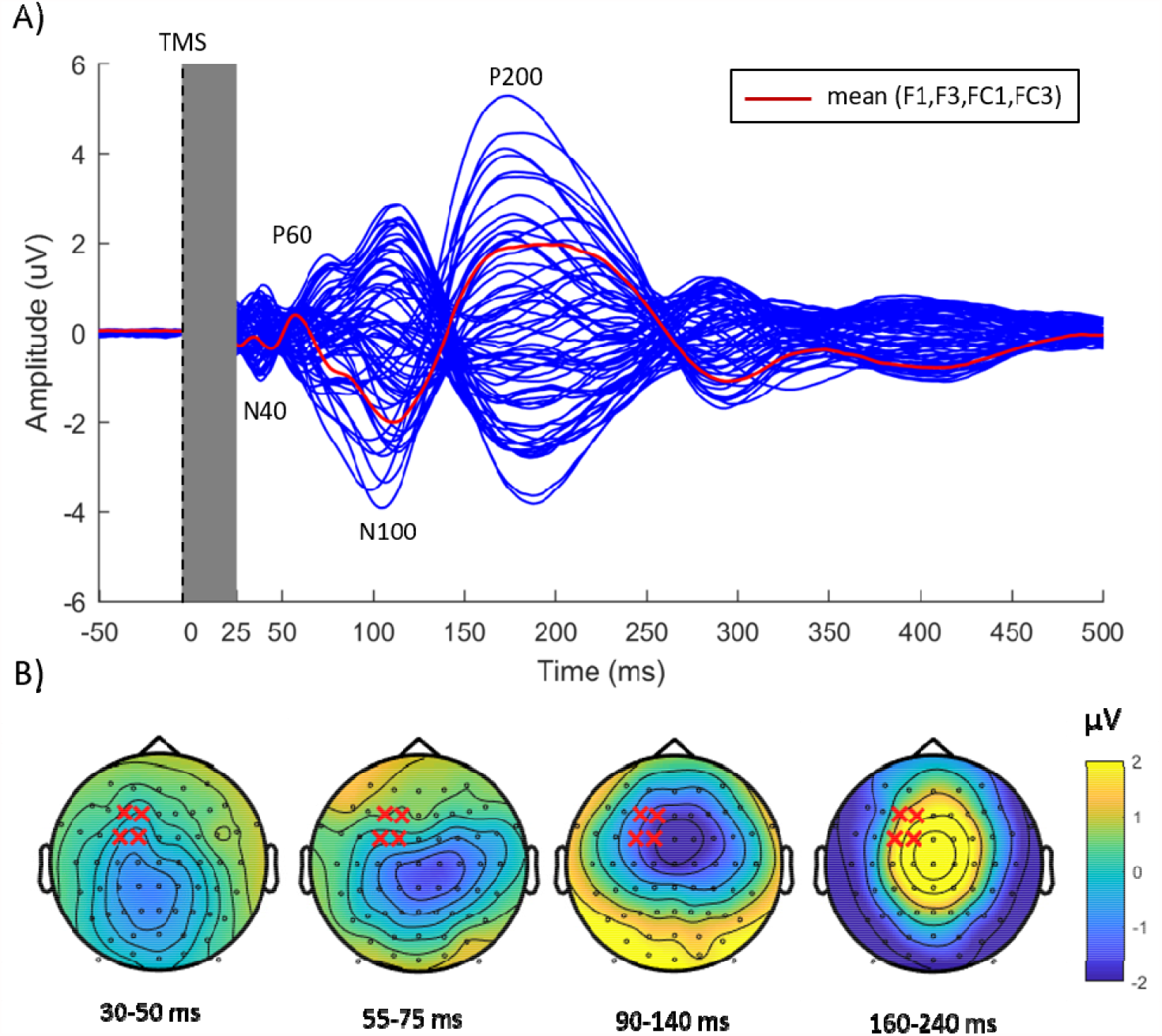
TMS-evoked potential following single-pulse stimulation over the left dorsolateral prefrontal cortex pre-TBS. TEPs were combined across the three different stimulation sessions and averaged across all participants. The grey box indicates removed data points due to TMS-related artifacts and cleaning steps. (**A**) Butterfly plot from all electrodes with major peaks (N40, P60, N100, P200) indicated in the text. The red lines indicate the waveform obtained from the mean of four electrodes (F3, FC3, F1, FC1) around the stimulation site. (**B**) Voltage distribution for each peak of interest averaged across time indicated below. The Red ‘x’ marks indicate the four electrodes chosen for the region of interest.

### 2.7 Statistical analysis

All statistical analyses were performed using either SPSS (version 26, IBM Corp, Armonk, NY, USA) or Matlab (version 2017b, MathWorks, Natick, MA, USA).

#### 2.7.1 ROI analysis

The local aftereffects in cortical reactivity of different forms of TBS were assessed in the predefined ROI over the left DLPFC (mean of F1, F3, FC1, FC3) using Mixed-Effects Repeated Measures Models (MRMMs). Fixed factors were Time (T2, T15 and T30), Condition (iTBS, cTBS, sham), and the Time × Condition interaction. ‘Participant’ was included as a random effect. The corresponding pre-TBS TEP amplitude (Pre) for each component was included as a covariate to account for individual differences in TEP amplitude at baseline. Restricted maximum likelihood estimation was used with an unstructured variance-covariance structure, which better fit most models (lowest Akaike’s Information Criterion (AIC)). For each component, amplitude values greater than three standard deviations from the grand mean of all experimental blocks together were considered outliers and excluded from analyses. Residuals were visually inspected to test for normality violations and ensure adequate convergence of model parameters (Schielzeth et al., 2020). Post-hoc pairwise comparisons were used to investigate significant main and interaction effects. Bootstrapping was applied to estimate mean effect sizes and 95% confidence intervals.

#### 2.7.2 Global scalp analysis

In addition to the predefined ROI analyses, we also performed data-driven exploratory analyses (covering the latency between 25 and 300 ms post-TMS) to assess changes in cortical reactivity across all electrodes on the scalp (Chung et al., 2017, Rogasch et al., 2020). To control for multiple comparisons across channels and time points, nonparametric, cluster-based permutation statistics were applied (Maris and Oostenveld, 2007) as implemented in Fieldtrip (https://www.fieldtriptoolbox.org/). The clusters were defined as two or more adjacent time samples or neighbouring EEG electrodes with a p-value < 0.05 in a dependent samples repeated-measures ANOVA. That is, to test for alterations in cortical reactivity following the different forms of TBS, difference scores at T2, T15 and T30 relative to Pre-TBS (Post-TBS at each of the three time points minus Pre-TBS: e.g., [iTBS_T15_ – iTBS_Pre_] vs [Sham_T15_ – Sham_Pre_]) were compared across conditions (iTBS, cTBS, sham). To test for differences in cortical reactivity over time within each stimulation condition, Post-TBS amplitudes at T2, T15 and T30 were compared to the Pre-TBS amplitude. Based on 5000 random permutations, Monte Carlo p-values were subsequently computed to form a distribution by means of a two-tailed test (p < 0.025).

#### 2.7.3 Signal-to-noise ratio (SNR)

The SNR was estimated for the predefined ROI for each participant individually by dividing the peak TEP amplitude of the component by the standard deviation of the waveform in the pre-stimulus interval (i.e. −500 to −50 ms), as used in previous TEP (Chung et al., 2017) and ERP studies (Debener et al., 2007, Hu et al., 2010). We also used the approach proposed by Luck et al. (2020) that estimates data quality using a ratio between the TEP amplitude (signal) and the standardised measurement error (SME) estimated by bootstrapping for that amplitude (noise).

## 3. Results

Data from 24 participants (72 sessions) were analysed (there were no dropouts). The average RMT was 65.5±7.6% of the maximum stimulator output (MSO). Overall, all interventions were well tolerated, with the most common side effect being a moderate headache (two cases during iTBS and one during cTBS; see Supplementary Material S2 for details).

### 3.1. Assessment of TEPs

The grand average of the TEP waveforms at the pre-TBS block is illustrated in Fig. 3A. Single-pulse TMS over the left DLPFC generated a series of deflections in the EEG traces, including two negative (N40, N100) and two positive (P60 and P200) components, classified according to their latencies after the TMS pulse (Chung et al., 2017, Chung et al., 2018b, Conde et al., 2019). Each component showed a distinctive topology (Fig. 3B), consistent with previous TMS-EEG studies in the prefrontal cortex (Chung et al., 2017, Hill et al., 2017, Conde et al., 2019). The latency and amplitude of each component were extracted for each participant and selected for further analysis (Supplementary Table S3). There was no significant difference in the latencies among the different stimulation conditions at the pre-TBS block (all p > 0.05).

For the ROI analyses, blocks with amplitudes greater than 3 SDs from the grand mean were considered outliers and excluded from analyses. A total of 3.4% of the data (10 out of 288 blocks) were excluded for the N40, 1.4% of data for P60 (4 out of 288 blocks), 5.5% for N100 (16 out of 288 blocks) and 5.9% for the P200 (17 out of 288 blocks). The final number of valid TEP trials per block included in the analysis (88 on average) is considered in the range for optimal TEP reliability following stimulation of the DLPFC (Kerwin et al., 2018). TEP waveforms were consistent across conditions and time points with distinct peaks in all time intervals of interest (Fig. 4).

**Fig. 4.**
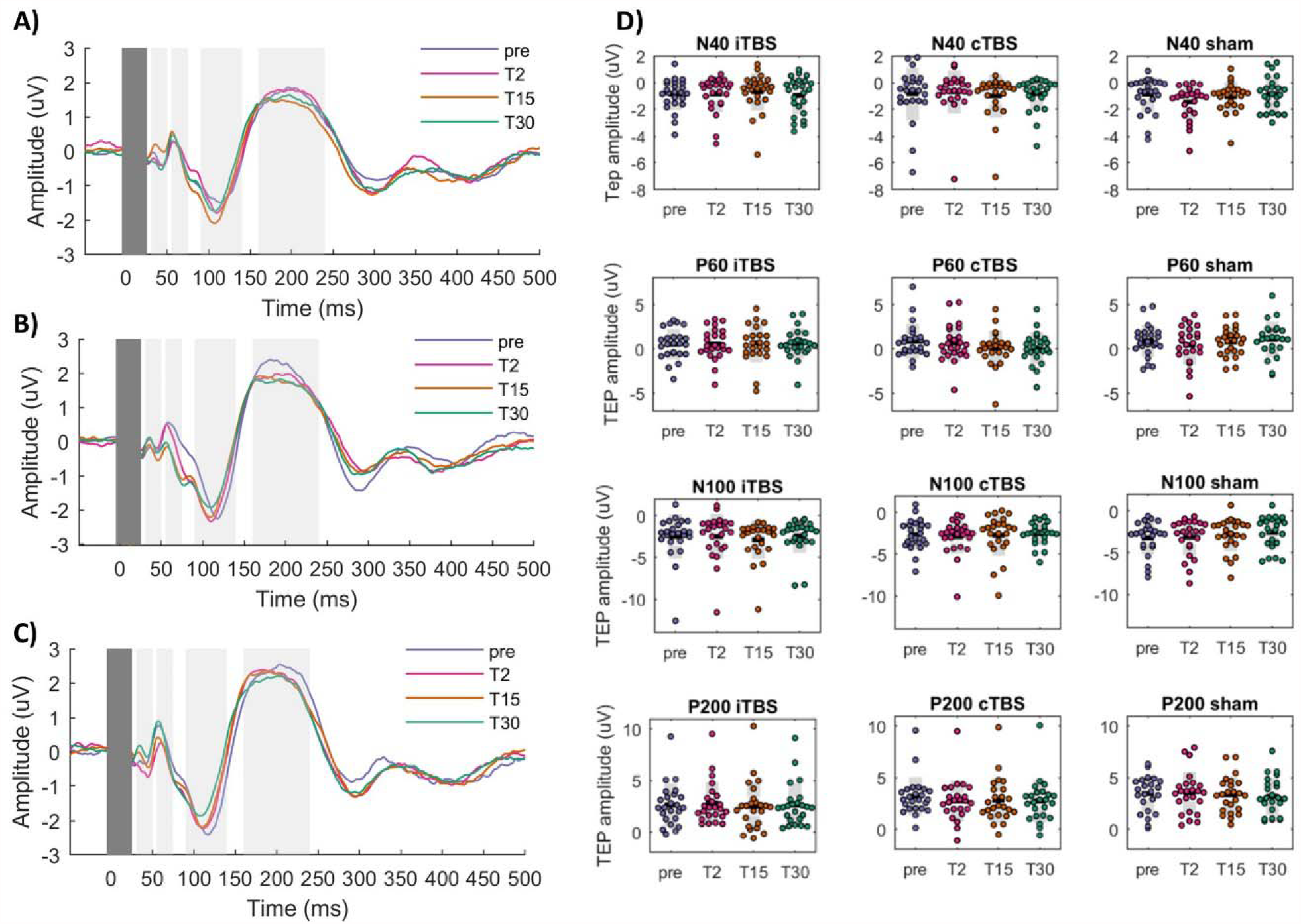
Transcranial magnetic stimulation (TMS)-evoked potentials (TEPs) before and after each stimulation condition for the average of F1, F3, FC1 and FC3. **(A)** Intermittent theta-burst stimulation (iTBS); **(B)** Continuous theta-burst stimulation (cTBS); **(C)** Sham. Grand average TEP waveforms pre-(blue) and post-TBS (2 min magenta, 15 min orange, 30 min green) for each stimulation conditions; **(D)** scatter plots showing the data of individual participants, the black lines show the mean, light grey shaded boxes indicate the standard deviation and dark grey regions indicate the 95% confidence interval.

We assessed the quality of the extracted TEPs using two approaches. Using the criteria proposed by Chung et al. (2017), all TEP components had SNR values above 3 SD, which was considered a reliable threshold for distinguishing signal from background noise, except for the N40 for the iTBS T2 block, cTBS T30 block and Sham pre-TBS block (Supplementary Table S4 and Fig. S4). Using the method proposed by Luck et al. (2020), only the SNR values for the P60 peak (at T2, T15, and T30 for iTBS and cTBS and T2 for sham) were below a recommended threshold of 10 (Luck, 2005) (Supplementary Table S5). Qualitatively, earlier peaks (N40 and P60) had a lower SNR than the later ones (N100 and P200). However, there was large inter-individual variability in TEP amplitude modulation’s magnitude and direction (increase/decrease) in all experimental conditions and time points (Fig. 4D and Supplementary Table S5, respectively).

### 3.2. ROI analysis of the effect of TBS on TEPs

MRMM analyses of the TEP amplitudes showed no significant main effects of Time, Condition, or Time x Condition interaction effect for any of the TEP components that were assessed (N40, P60, N100, P200; Table 1). Only the N40 Condition effect (F(2,14.4) = 3.5, p = 0.056) and the N100 interaction effect (F(4, 14.7) = 2.6, p = 0.078) were close to the significance threshold. The estimated effect sizes between conditions ranged from small to large, with wide confidence intervals (Supplementary Material, Fig. S6 and Table S6). The strongest effect relative to sham was found with iTBS at T15 for the N100 (Cohen’s d = −0.5, 95%CI −1.14 to 0.16). When all conditions were compared, the strongest effect size was also found for the N100 component at T15 between iTBS and cTBS (Cohen’s d = −0.76, 95%CI - 1.39 to −0.14), with N100 amplitudes being more negative for iTBS than with cTBS. Likewise, the effect sizes of the pre-post differences in TEPs were small (Supplementary Material, Fig. S7 and Table S7). For the N100, the effect size relative to baseline was greatest at T15 for both active conditions, but in opposite directions; more negative amplitudes after iTBS (Cohen’s d = −0.3, 95%CI −0.9 to 0.33) and less negative after cTBS (Cohen’s d = 0.33, 95%CI - 0.26 to 1.04). The effects of sham stimulation in N100 amplitudes were negligible throughout the session (all effect sizes < 0.04).

**Table 1.** Mixed Effects Repeated Measures Model (MRMM) for the ROI analysis of the effect of TBS on TEPs

|  |  | df1 | df2 | F | p |
| --- | --- | --- | --- | --- | --- |
| N40 | Time | 2 | 17.1 | 0.794 | 0.470 |
|  | Condition | 2 | 14.4 | 3.546 | 0.056 |
|  | Time x Condition | 4 | 21.5 | 1.386 | 0.272 |
| P60 | Time | 2 | 21.3 | 0.111 | 0.895 |
|  | Condition | 2 | 22.4 | 0.990 | 0.387 |
|  | Time x Condition | 4 | 22.0 | 1.288 | 0.305 |
| N100 | Time | 2 | 15.15 | 0.104 | 0.902 |
|  | Condition | 2 | 14.9 | 0.192 | 0.827 |
|  | Time x Condition | 4 | 14.7 | 2.611 | 0.078 |
| P200 | Time | 2 | 18.2 | 0.576 | 0.572 |
|  | Condition | 2 | 16.0 | 2.639 | 0.102 |
|  | Time x Condition | 4 | 17.8 | 1.053 | 0.408 |

### 3.3 Global scalp analyses

In addition to our ROI analyses, we also performed data-driven exploratory analyses to assess differences in TEPs across all EEG channels. Using cluster-based permutation tests, we did not find significant differences in TEPs between conditions for any of the time points (all p > 0.025, two-tailed test), in line with our ROI analysis.

In addition, we performed permutation analyses to investigate changes from baseline to the post-TBS blocks for sham, cTBS, and iTBS, the results of which are reported in Supplementary Material S10.

## Discussion

This study examined the electrophysiological effects of TBS applied to the left DLPFC using TMS-evoked potentials (TEPs). The amplitudes of the N40, P60, N100 and P200 components were assessed before and after iTBS, cTBS and sham. Contrary to our expectations, we did not find any significant differences in the N100 amplitude between stimulation conditions at our ROI (average of channels F1, F3, FC1 and FC3). This was confirmed by the cluster-based permutation tests, which did not detect any clusters that revealed significant differences between conditions.

In contrast, the original TMS-EEG study in 10 healthy volunteers on which the current experiment was based showed a significant increase in N100 amplitude between iTBS and sham and after iTBS compared to baseline in the same ROI, although there was no difference with cTBS (Chung et al., 2017). Our MRMM results showed an interaction effect on the N100 close to the significance threshold, with the strongest effect at T15, similarly indicating an increased N100 amplitude post iTBS compared to sham. Despite not being statistically significant, this difference was in the same direction as hypothesised, consistent with previous sham-controlled TBS studies (Chung et al., 2017, Chung et al., 2018a). Although the underlying electrophysiology of TEPs has not been completely elucidated, the N100 component is thought to be related to inhibitory GABA-ergic activity both in the motor (Premoli et al., 2014) and prefrontal regions (Rogasch et al., 2015).

Some methodological differences may explain why we achieved a smaller effect size and could not detect a significant difference between conditions in the local TEPs compared to the study we used to base our conceptual replication. The intensity for the active TBS conditions used in the present experiment was 75% compared to 80% RMT in the original study (Chung et al., 2017). This modification was adopted following evidence that the strongest effect on cortical reactivity in the prefrontal cortex was produced using a slightly weaker iTBS intensity of 75% RMT (Chung et al., 2018b), as well to increase tolerability by minimising participant discomfort. However, considering the large inter-individual variability in response to TBS (Chung et al., 2017, Chung et al., 2019), this intensity may not have been ideal for our sample, potentially being too weak to achieve reliable neural modulation in all participants and thereby mitigating TBS effects.

The waveform of single TMS pulses used to evoke TEPs was also different between experiments (biphasic in the present versus monophasic in the original), which may also have contributed to the discrepancy in findings. Pulse shapes affect neural populations differently (Casula et al., 2018, Pisoni et al., 2018) and result in different artifact profiles (Rogasch et al., 2013, Huang and Mouraux, 2015), even with stimulus intensity normalised to individual RMT. For example, biphasic pulses delivered to the premotor cortex evoked larger early TEP components (e.g. around 40 ms, i.e. N40), whereas monophasic pulses elicited greater middle-latency components (e.g. around 100 ms, i.e. N100) (Pisoni et al., 2018).

Although the current study was powered to detect a moderate-to-large effect size (Cohen’s d = −0.60), the largest effect sizes on the change of N100 was Cohen’s d =-0.50, smaller than we were powered to detect and far smaller than the original study, which examined pre to post TBS changes (Cohen’s d=-1.50). The sample size of the original study was ten subjects, and effect size estimates are generally inflated in small samples (Szucs and Ioannidis, 2020). For comparison, a meta-analysis of the effect of iTBS on motor evoked potentials (MEPs) found a significant increase in MEP at early time points (within the first 5 min) with a corrected pooled standardised mean difference (SMD) of 0.36 (Chung et al., 2016). In the current study, the largest effect sizes with iTBS in the change of N100 amplitude from baseline was found at the T15 block (Cohen’s d = −0.30), smaller than the original study (Cohen’s d=-1.04) and closer to the estimated effect size on motor cortical excitability. The large confidence intervals indicate large inter-individual variability in response to TBS, suggesting that the heterogeneity of TBS effects at the prefrontal cortex is similar to those found in TBS experiments conducted at the motor cortex (Corp et al., 2020, Ozdemir et al., 2021). Therefore, a more reliable estimate of the population effect size of TBS administered to the DLPFC may require studies with larger sample sizes, meta-analyses integrating results from multiple datasets or Bayesian approaches.

An assessment of the local effects of TBS via a predefined ROI over the DLPFC was only performed in one study (Chung et al., 2017) and disregarded in subsequent TBS experiments in favour of a global scalp exploratory approach using nonparametric cluster-based permutation statistics (Chung et al., 2018a, Chung et al., 2018b, Chung et al., 2019). Even with corrections for multiple comparisons, global scalp cluster-based approaches may inflate false-positive rates, although the inflation size is debated (Cox et al., 2017). Restriction of analyses within specific brain areas, as with the use of a pre-registered ROI, offers a better balance between sensitivity and the number of performed comparisons (Gentili et al., 2021). Alternatively, the use of a collapsed localiser approach (Luck and Gaspelin, 2017, Cohen, 2014), blind to group condition, to select the region of interest may provide an intermediary compromise between preregistered ROIs and exploratory data-driven channel selection. As more data is published on the topic and larger datasets become available, these can be leveraged to determine the appropriate ROI to assess the local effects of TBS and, thereby, improve the validity of NIBS techniques that target the prefrontal cortex. Therefore, future studies are needed to determine the appropriate ROI to assess the local effects of TBS and improve the validity of NIBS techniques that target prefrontal areas in clinical settings.

## Limitations

There are some limitations of this study that must be acknowledged. A spacer was used to minimise the introduction of artifacts in the EEG electrodes under the Magventure coil, arising due to electrode movement and bone-conducted auditory input. This procedure increased the distance between the coil and the scalp, requiring relatively higher stimulation intensities and yielding a louder TMS clicking sound. Even with white-noise auditory masking and earmuffs, this larger auditory component was only partially mitigated. Our experimental design did not include a sensory control TEP condition using a form of “realistic sham” mimicking the auditory and somatosensory co-stimulation (Gordon et al., 2018, Conde et al., 2019). However, we used the same stimulation intensity during all sessions, ensuring that the contribution of peripheral auditory and somatosensory evoked potentials to the TEP measurement remained constant. Therefore, it is expected that these peripheral artifacts were consistent across time and that changes in TEP amplitudes reflected changes in neural excitability due to TMS. Another potential limitation is that the localisation of DLPFC for the TMS coil was based on the F3 electrode location and not stereotaxic neuronavigation. Although electrode-based positioning is commonly employed in TMS-EEG research (Chung et al., 2017, Cash et al., 2017, Chung et al., 2018b), individualised targeting of the DLPFC could be improved by using MRI-guided neuronavigation. Neuronavigation further allows real-time monitoring of coil position and angle relative to scalp placement, which has recently been advocated to reduce variability in TEP measurements (Varone et al., 2021).

## Conclusion

The present study used TMS-evoked potentials (TEPs) to assess TBS’s neuromodulation effects at the prefrontal cortex. We did not find significant differences in cortical reactivity between iTBS, cTBS and sham. The pooled effect size of N100 amplitudes following iTBS was smaller than previously reported in the literature, suggesting that the true effect size may be more modest, similar to aggregate effect size estimates for stimulation of the motor cortex. Larger samples, meta-analytic methods, or Bayesian approaches using informative priors are needed to overcome inter-individual heterogeneity and accurately assess TBS effects at the prefrontal cortex, potentially uncovering moderating factors for the variability in responses. To support these future endeavours, the data used in the present study will be presented in conjunction with EEG pre-processing scripts in an upcoming data paper.

## Acknowledgements

The authors wish to thank Nichola Jephcott, Design Futures Lab, School of Built Environment, UNSW, for the assistance with the customised 3D printed spacer, and all volunteers for their participation. A.H. Moffa is the recipient of a Scientia PhD Scholarship from the University of New South Wales, Sydney, Australia. The present work will contribute to the doctoral thesis of A. H. Moffa.

## Notes

### Competing Interest Statement

The authors have declared no competing interest.

### Summary of Updates

Figure 1 updated due to poor resolution Figure 3 revised due to missing subtitle

