## Supplementary Material for "Assessing neuromodulation effects of theta burst stimulation to the prefrontal cortex using TMS-evoked potentials" (from another version of this preprint: version 1, 2021-06-22)

S1. Resting motor threshold (RMT)

Resting motor threshold (RMT) was measured using the procedure detailed in (Conforto et al., 2004). Briefly, MEPs were assessed using Ag/AgCl EMG electrodes placed over the first dorsal interosseous (FDI) muscle. The hot spot was identified, and the individual RMTs at the scalp and with the spacer were defined as the lowest intensity (the percentage of the maximum stimulator output, MSO) necessary to elicit MEPs with a peak-to-peak amplitude of > 0.05mV in at least 3 out of 6 trials (Conforto et al., 2004).

S2. Adverse effects of TBS

Two participants reported moderate headache and one mild nausea during iTBS. One participant reported mild blurred vision, one participant moderate headache and fatigue, and one mild nausea and moderate dizziness during cTBS. One participant reported moderate fatigue during sham stimulation. All other participants underwent the measurements without reporting any other adverse effects.

S3. Latencies of peaks

**Table S3:** Latencies of each peak at baseline (pre) and at the three time points after each stimulation condition (M±SD).

| **iTBS** | Pre | T2 | T15 | Post3 |
| --- | --- | --- | --- | --- |
| N40 | 39.9 ± 8.3 | 39.7 ± 8.8 | 40.0 ± 8.4 | 38.8 ± 7.7 |
| P60 | 61.6 ± 7.2 | 62.9 ± 6.5 | 61.7 ± 7.3 | 59.4 ± 6.2 |
| N100 | 119.9 ± 15.7 | 113.1 ± 14.8 | 109.1 ± 13.7 | 111.5 ± 13.6 |
| P200 | 204.9 ± 25.2 | 200.5 ± 26.6 | 191.9 ± 26.5 | 201.0 ± 29.1 |

| **cTBS** | Pre | T2 | T15 | Post3 |
| --- | --- | --- | --- | --- |
| N40 | 39.2 ± 7.8 | 40.9 ± 7.9 | 40.7 ± 7.9 | 39.9 ± 8.4 |
| P60 | 61.5 ± 7.1 | 59.1 ± 5.7 | 61.9 ± 8.0 | 58.8 ± 6.0 |
| N100 | 115.5 ± 11.6 | 110.6 ± 11.16 | 108.9 ± 13.1 | 109.7 ± 12.7 |
| P200 | 195.7 ± 27.3 | 194.7 ± 25.2 | 187.8 ± 22.9 | 191.1 ± 27.4 |

| **Sham** | Pre | T2 | T15 | Post3 |
| --- | --- | --- | --- | --- |
| N40 | 37.2± 6.9 | 39.2±7.9 | 40.7 ± 7.8 | 40.9 ± 7.7 |
| P60 | 59.4± 4.7 | 61.2±5.2 | 60.7 ± 6.2 | 59.4 ± 4.9 |
| N100 | 115.5± 15.4 | 109.8±13.6 | 111.7 ± 15.8 | 109.0 ± 15.3 |
| P200 | 201.3± 24.9 | 194.1±24.6 | 200.4 ± 26.2 | 194.2 ± 26.2 |

S4. Signal to noise ratio (SNR)

**Table S4** summarises the averaged SNR of single-pulse TMS included in the analyses. The values represent how many standard deviations each peak amplitude was above baseline. A value of 3 SDs (99.7% of the data) or more was considered a good SNR. Qualitatively N40 and P60 peaks displayed a moderate-to-good SNR, but N100 and P200 exhibited excellent SNR.

| **iTBS** | Pre | T2 | T15 | Post3 |
| --- | --- | --- | --- | --- |
| N40 | 3.1±3.3 | 2.8±3.3 | 3.0±4.6 | 3.5±3.9 |
| P60 | 3.5±2.6 | 3.9±3.7 | 4.5±4.6 | 3.5±3.5 |
| N100 | 7.4±8.8 | 7.9±7.9 | 8.5±9.0 | 6.2±4.2 |
| P200 | 7.1±6.7 | 7.8±5.8 | 7.4±8.3 | 7.0±4.9 |

| **cTBS** | Pre | T2 | T15 | Post3 |
| --- | --- | --- | --- | --- |
| N40 | 4.0±4.0 | 3.1±2.9 | 3.0±2.9 | 2.5±2.5 |
| P60 | 4.2±5.0 | 4.5±4.1 | 3.4±3.2 | 3.1±2.8 |
| N100 | 8.1±6.0 | 9.3±6.3 | 7.8±6.1 | 7.4±4.4 |
| P200 | 9.0±4.9 | 9.0±6.5 | 8.4±6.1 | 7.7±4.5 |

| **Sham** | Pre | T2 | T15 | Post3 |
| --- | --- | --- | --- | --- |
| N40 | 2.9±3.1 | 4.4±4.3 | 3.2±2.4 | 3.8±3.4 |
| P60 | 3.56±2.4 | 4.4±3.7 | 4.1±4.0 | 4.9±4.9 |
| N100 | 8.5±6.3 | 8.6±7.0 | 7.9±7.2 | 7.2±6.4 |
| P200 | 8.6±5.5 | 9.0±5.7.0 | 8.6±5.7 | 8.4±6.8 |

**Figure S4.** Signal-to-noise ratio for each experimental condition in the ROI (F1, F3, FC1, FC3). The SNR was estimated for the predefined ROI for each participant individually by dividing the peak TEP amplitude of the component by the standard deviation of the waveform in the pre-stimulus interval (i.e., -500 to -50 ms). A) N40; B) P60; C) N100; D) P200.

A) N40

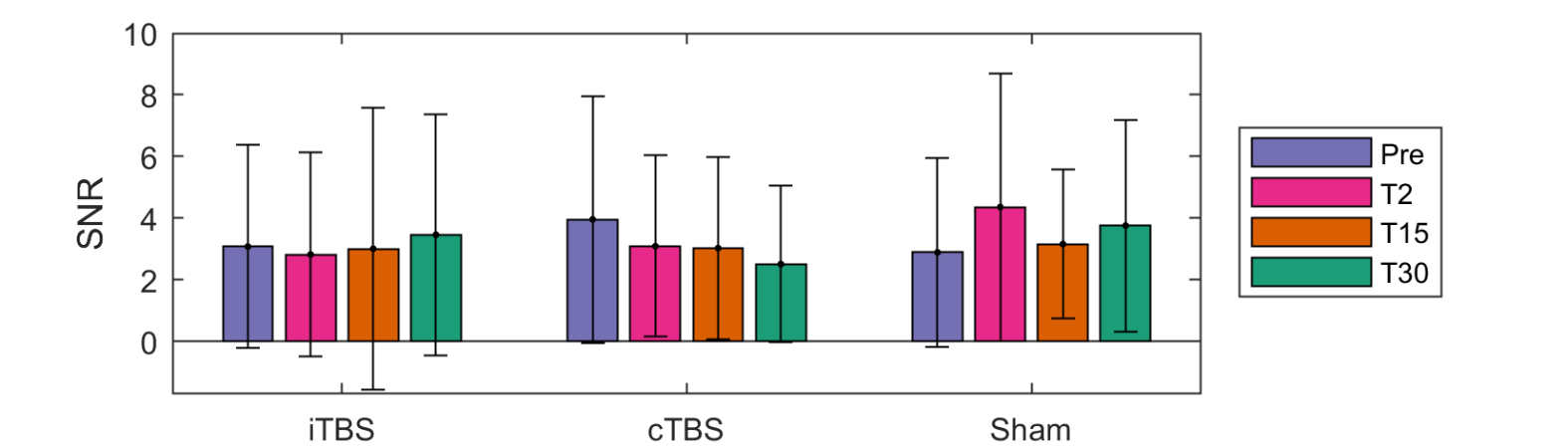

B) P60

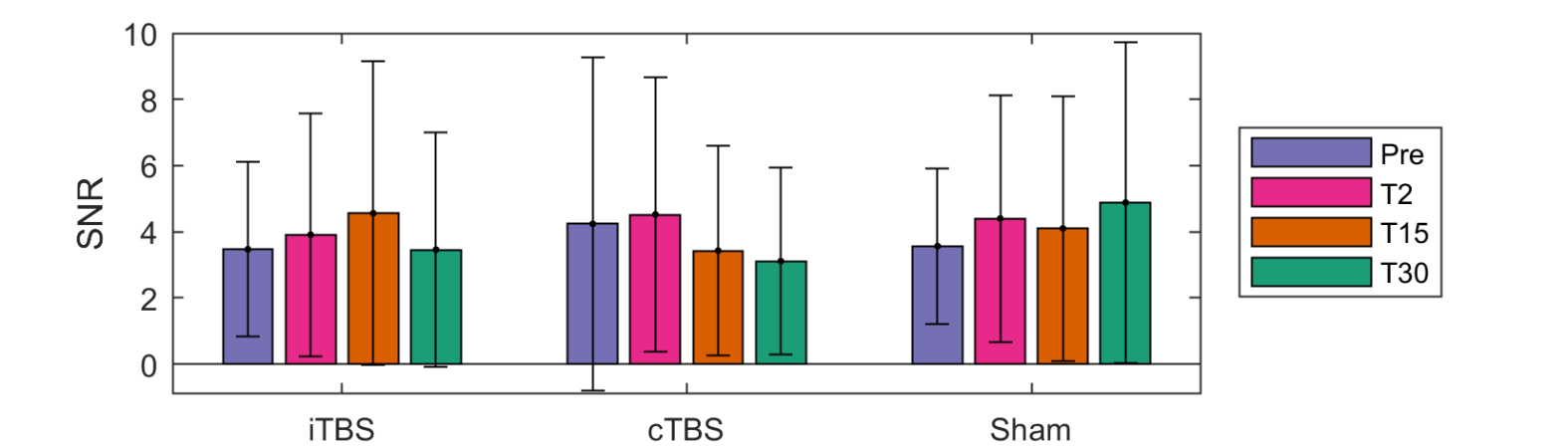

C) N100

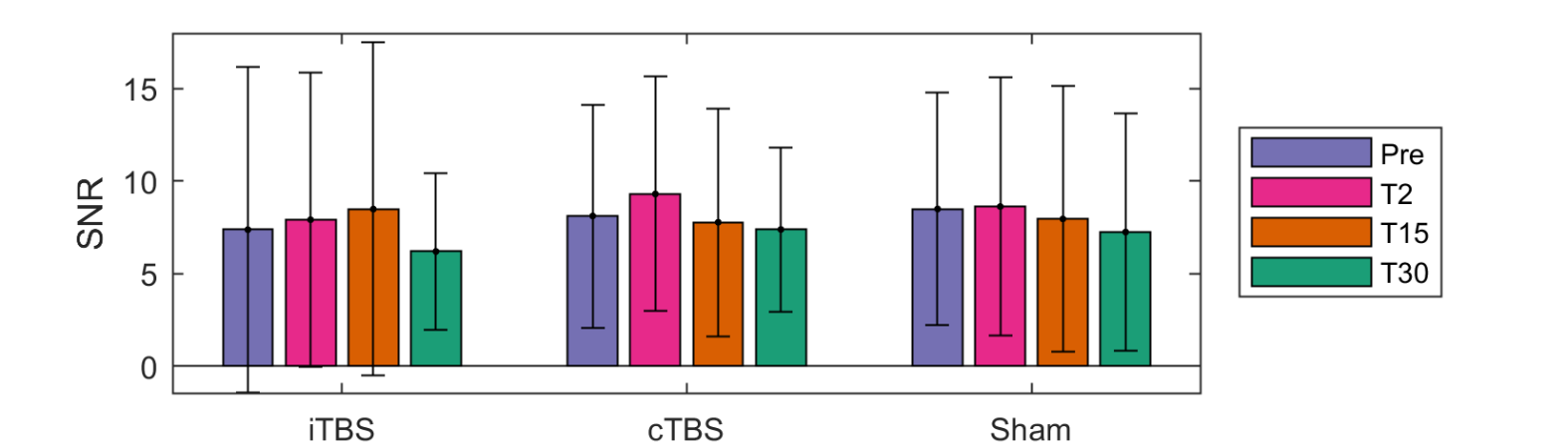

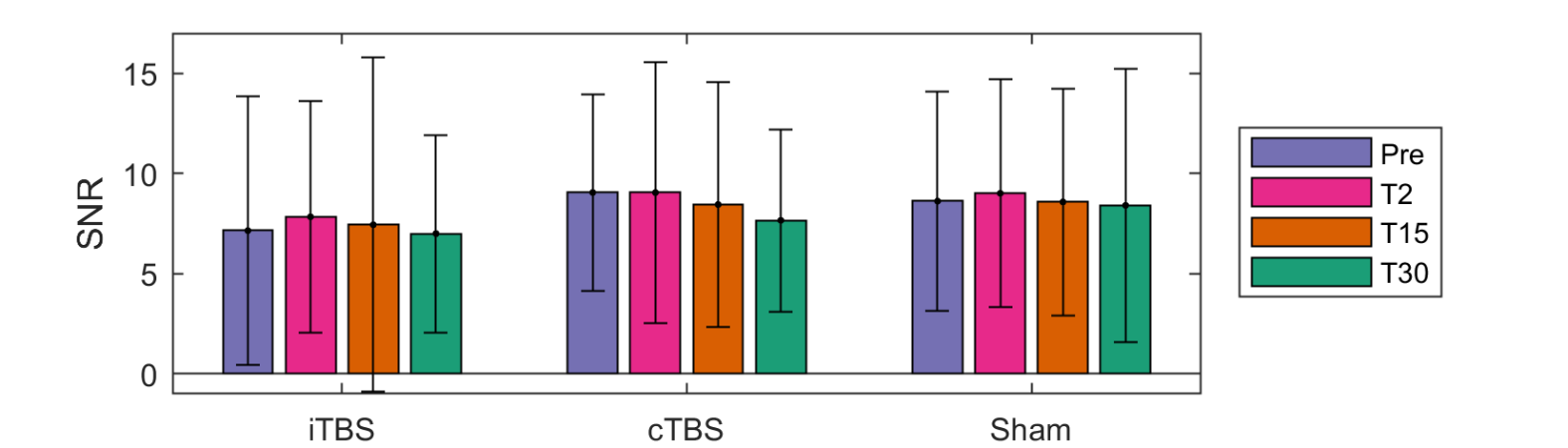
D) P200

**Table S5** summarises the averaged SNR of single-pulse TMS included in the analyses using Luck et al. 2020. The values represent how many standard deviations each peak amplitude was above baseline. A SNR of 10 is the recommended threshold for ERP studies (Luck, 2005).

| **iTBS** | Pre | T2 | T15 | Post3 |
| --- | --- | --- | --- | --- |
| N40 | 18.8 | 16.4 | 11.9 | 17.1 |
| P60 | 10.6 | 7.7 | 7.6 | 8.9 |
| N100 | 22.8 | 23.8 | 30.5 | 29.6 |
| P200 | 31.2 | 33.2 | 25.3 | 30.4 |

| **cTBS** | Pre | T2 | T15 | Post3 |
| --- | --- | --- | --- | --- |
| N40 | 11.0 | 10.8 | 15.7 | 17.2 |
| P60 | 11.2 | 7.6 | 0.7 | 1.2 |
| N100 | 36.2 | 36.8 | 27.0 | 44.4 |
| P200 | 42.8 | 33.8 | 32.3 | 33.6 |

| **Sham** | Pre | T2 | T15 | Post3 |
| --- | --- | --- | --- | --- |
| N40 | 17.0 | 28.7 | 23.4 | 16.3 |
| P60 | 16.6 | 4.7 | 11.3 | 13.4 |
| N100 | 40.3 | 36.3 | 36.1 | 32.1 |
| P200 | 49.0 | 43.0 | 46.8 | 43.1 |

S5. Percentage of subjects in which TEPs increased/decreased

**Table S5**. Percentage of subjects in which TEPs increased/decreased relative to pre-stimulation (complete data with no outliers excluded).

|  | | **N40** | | | **P60** | | | **N100** | | | | **P200** | | |
| --- | --- | --- | --- | --- | --- | --- | --- | --- | --- | --- | --- | --- | --- | --- |
|  |  | **iTBS** | **cTBS** | **Sham** | **iTBS** | **cTBS** | **Sham** | **iTBS** | **cTBS** | **Sham** | **iTBS** | | **cTBS** | **Sham** |
|  |  | 62.5% | 50.0% | 33.3% | 45.8% | 54.2% | 41.7% | 54.2% | 45.8% | 54.2% | 70.8% | | 41.7% | 62.5% |
| **T2** |  |  |  |  |  |  |  |  |  |  |  |  |  |  |
|  |  | 37.5% | 50.0% | 66.7% | 54.2% | 45.8% | 58.3% | 45.8% | 54.2% | 45.8% | 29.2% | | 58.3% | 37.5% |
|  |  | 58.3% | 41.7% | 29.2% | 54.2% | 33.3% | 37.5% | 33.3% | 54.2% | 50.0% | 62.5% | | 33.3% | 50.0% |
| **T15** |  |  |  |  |  |  |  |  |  |  |  |  |  |  |
|  |  | 41.7% | 58.3% | 70.8% | 45.8% | 66.7% | 62.5% | 66.7% | 45.8% | 50.0% | 37.5% | | 66.7% | 50.0% |
| **T30** |  | 50.0% | 45.8% | 41.7% | 54.2% | 33.3% | 50.0% | 58.3% | 54.2% | 62.5% | 45.8% | | 45.8% | 41.7% |
|  |  | 50.0% | 54.2% | 58.3% | 45.8% | 66.7% | 50.0% | 41.7% | 45.8% | 37.5% | 54.2% | | 54.2% | 58.3% |

S6. Effect sizes between conditions in the predefined ROI (F1, F3, FC1, FC3)

**Table S6.** Effect sizes- between conditions of the relative change between the pre-stimulation block and the three post time points. Bootstrapping was applied to estimate the median effect sizes (ES) and 95% confidence intervals (blocks with amplitude values greater than three standard deviations from the grand mean of all experimental blocks were considered outliers and excluded from analyses).

| **N40** | **iTBS vs. sham** | | | **cTBS vs. sham** | | | **iTBS vs. cTBS** | | |
| --- | --- | --- | --- | --- | --- | --- | --- | --- | --- |
| **Cohen's d** | **Mean** | **lb** | **ub** | **Mean** | **lb** | **ub** | **Mean** | **lb** | **ub** |
| **'T2'** | 0.41 | -0.22 | 1.11 | 0.37 | -0.31 | 1.01 | -0.02 | -0.65 | 0.61 |
| **'T15'** | 0.49 | -0.15 | 1.32 | 0.08 | -0.46 | 0.81 | 0.52 | -0.13 | 1.11 |
| **'T30'** | -0.06 | -0.62 | 0.53 | -0.19 | -0.78 | 0.39 | 0.18 | -0.37 | 0.91 |

| **P60** | **iTBS vs. sham** | | | **cTBS vs. sham** | | | **iTBS vs. cTBS** | | |
| --- | --- | --- | --- | --- | --- | --- | --- | --- | --- |
| **Cohen's d** | **Mean** | **lb** | **ub** | **Mean** | **lb** | **ub** | **Mean** | **lb** | **ub** |
| **'T2'** | 0.19 | -0.41 | 0.74 | 0.05 | -0.53 | 0.69 | 0.12 | -0.53 | 0.71 |
| **'T15'** | 0.20 | -0.36 | 0.85 | -0.26 | -0.85 | 0.27 | 0.50 | -0.11 | 1.17 |
| **'T30'** | -0.02 | -0.62 | 0.52 | -0.48 | -1.13 | 0.10 | 0.48 | -0.10 | 1.10 |

| **N100** | **iTBS vs. sham** | | | **cTBS vs. sham** | | | **iTBS vs. cTBS** | | |
| --- | --- | --- | --- | --- | --- | --- | --- | --- | --- |
| **Cohen's d** | **Mean** | **lb** | **ub** | **Mean** | **lb** | **ub** | **Mean** | **lb** | **ub** |
| **'T2'** | -0.09 | -0.82 | 0.49 | -0.11 | -0.88 | 0.48 | 0.04 | -0.58 | 0.61 |
| **'T15'** | -0.50 | -1.14 | 0.16 | 0.11 | -0.51 | 0.76 | -0.76 | -1.39 | -0.14 |
| **'T30'** | -0.20 | -0.85 | 0.38 | -0.19 | -0.84 | 0.37 | 0.03 | -0.54 | 0.68 |

| **P200** | **iTBS vs. sham** | | | **cTBS vs. sham** | | | **iTBS vs. cTBS** | | |
| --- | --- | --- | --- | --- | --- | --- | --- | --- | --- |
| **Cohen's d** | **Mean** | **lb** | **ub** | **Mean** | **lb** | **ub** | **Mean** | **lb** | **ub** |
| **'T2'** | 0.02 | -0.60 | 0.77 | -0.44 | -1.14 | 0.22 | 0.37 | -0.22 | 1.15 |
| **'T15'** | 0.06 | -0.54 | 0.70 | -0.17 | -0.85 | 0.43 | 0.17 | -0.42 | 0.83 |
| **'T30'** | 0.29 | -0.33 | 0.90 | 0.05 | -0.64 | 0.65 | 0.23 | -0.39 | 0.85 |

**Figure S****6**. Effect sizes of the relative change from baseline between A) iTBS vs sham, B) cTBS vs sham, C) iTBS vs cTBS.

A) iTBS vs sham

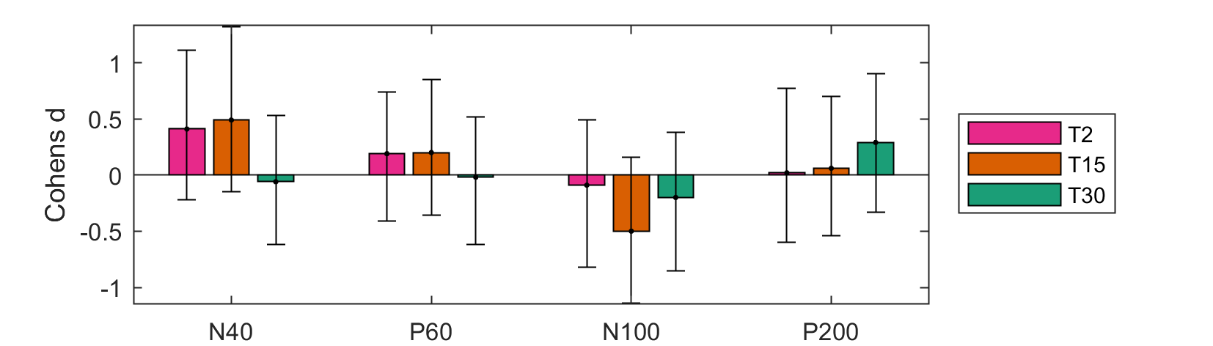

B) cTBS vs sham

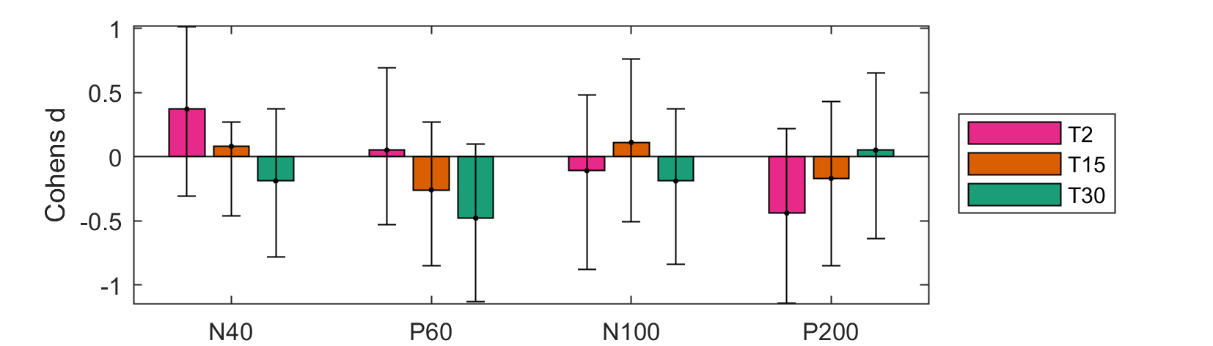

C) iTBS vs cTBS

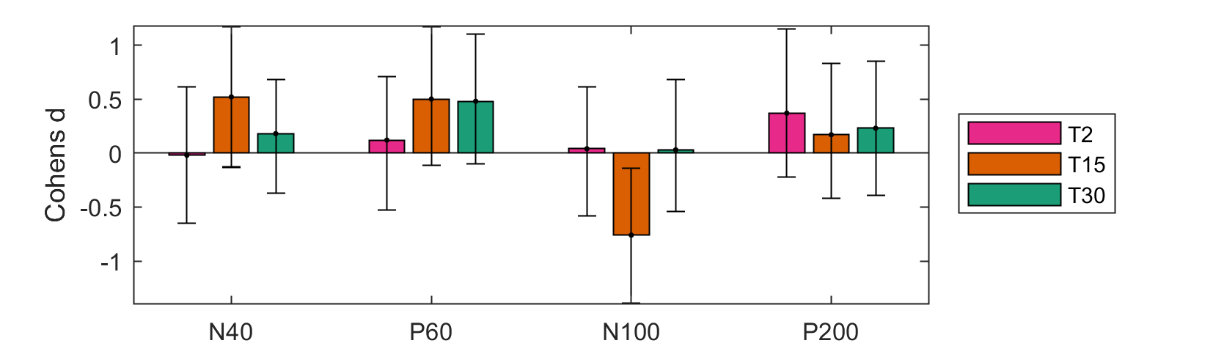

S7. Effect sizes within conditions in the predefined ROI (F1, F3, FC1, FC3)

**Table S7.** Effect sizes- within condition of the relative change from the pre-stimulation block across time. Bootstrapping was applied to estimate the median effect sizes (ES) and 95% confidence intervals (blocks with amplitude values greater than three standard deviations from the grand mean of all experimental blocks were considered outliers and excluded from analyses).

| **N40** | **iTBS** | | | **cTBS** | | | **Sham** | | |
| --- | --- | --- | --- | --- | --- | --- | --- | --- | --- |
| **Cohen's d** | **Mean** | **lb** | **ub** | **Mean** | **lb** | **ub** | **Mean** | **lb** | **ub** |
| **'T2 vs baseline'** | 0.01 | -0.54 | 0.61 | -0.04 | -0.58 | 0.59 | -0.40 | -1.12 | 0.24 |
| **'T15 vs baseline'** | 0.38 | -0.23 | 0.99 | -0.22 | -0.77 | 0.37 | -0.31 | -0.98 | 0.34 |
| **'T30 vs baseline'** | -0.07 | -0.64 | 0.48 | -0.28 | -0.82 | 0.31 | -0.11 | -0.74 | 0.50 |

| **P60** | **iTBS** | | | **cTBS** | | | **Sham** | | |
| --- | --- | --- | --- | --- | --- | --- | --- | --- | --- |
| **Cohen's d** | **Mean** | **lb** | **ub** | **Mean** | **lb** | **ub** | **Mean** | **lb** | **ub** |
| **'T2 vs baseline'** | 0.03 | -0.53 | 0.61 | -0.11 | -0.68 | 0.51 | -0.13 | -0.70 | 0.53 |
| **'T15 vs baseline'** | 0.10 | -0.44 | 0.66 | -0.19 | -0.82 | 0.37 | -0.10 | -0.67 | 0.51 |
| **'T30 vs baseline'** | 0.08 | -0.51 | 0.68 | -0.34 | -0.93 | 0.30 | 0.08 | -0.52 | 0.66 |

| **N100** | **iTBS** | | | **cTBS** | | | **Sham** | | |
| --- | --- | --- | --- | --- | --- | --- | --- | --- | --- |
| **Cohen's d** | **Mean** | **lb** | **ub** | **Mean** | **lb** | **ub** | **Mean** | **lb** | **ub** |
| **'T2 vs baseline'** | 0.12 | -0.46 | 0.70 | -0.09 | -0.71 | 0.53 | -0.03 | -0.62 | 0.59 |
| **'T15 vs baseline'** | -0.30 | -0.90 | 0.33 | 0.33 | -0.26 | 1.04 | 0.03 | -0.59 | 0.66 |
| **'T30 vs baseline'** | 0.14 | -0.47 | 0.74 | -0.05 | -0.64 | 0.56 | 0.04 | -0.54 | 0.70 |

| **P200** | **iTBS** | | | **cTBS** | | | **Sham** | | |
| --- | --- | --- | --- | --- | --- | --- | --- | --- | --- |
| **Cohen's d** | **Mean** | **lb** | **ub** | **Mean** | **lb** | **ub** | **Mean** | **lb** | **ub** |
| **'T2 vs baseline'** | -0.02 | -0.68 | 0.61 | -0.26 | -0.83 | 0.33 | -0.18 | -0.83 | 0.44 |
| **'T15 vs baseline'** | -0.14 | -0.72 | 0.46 | -0.20 | -0.88 | 0.43 | -0.14 | -0.81 | 0.46 |
| **'T30 vs baseline'** | -0.01 | -0.65 | 0.61 | -0.29 | -0.94 | 0.35 | -0.14 | -0.82 | 0.44 |

**Figure S7.** Effect sizes within each condition across time in relation to baseline A) iTBS, B) cTBS and C) sham. Outliers over 3 SD were excluded.

A) iTBS

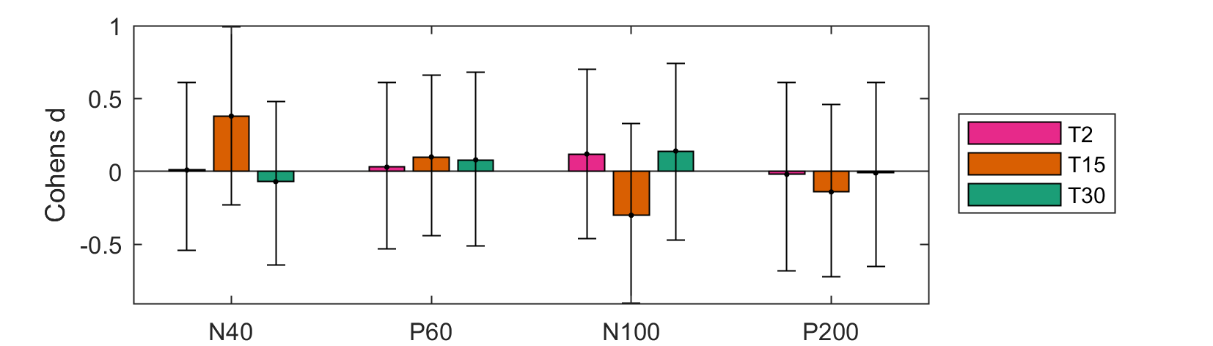

B) cTBS

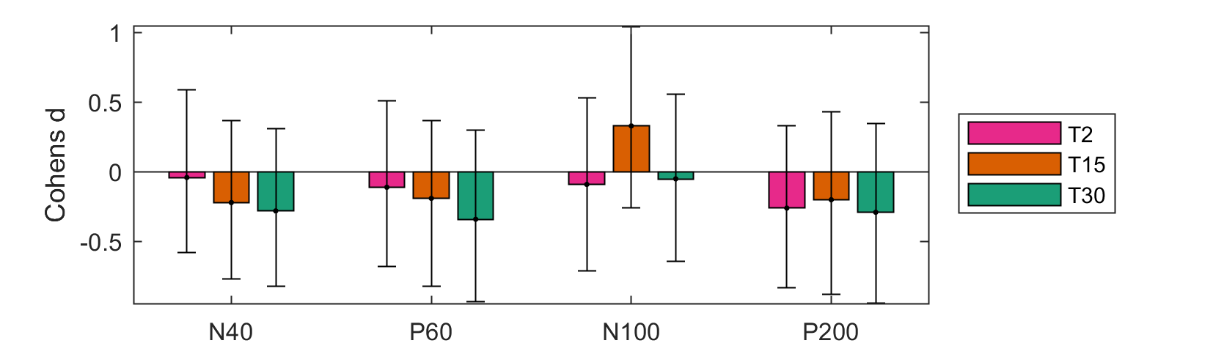

C) Sham

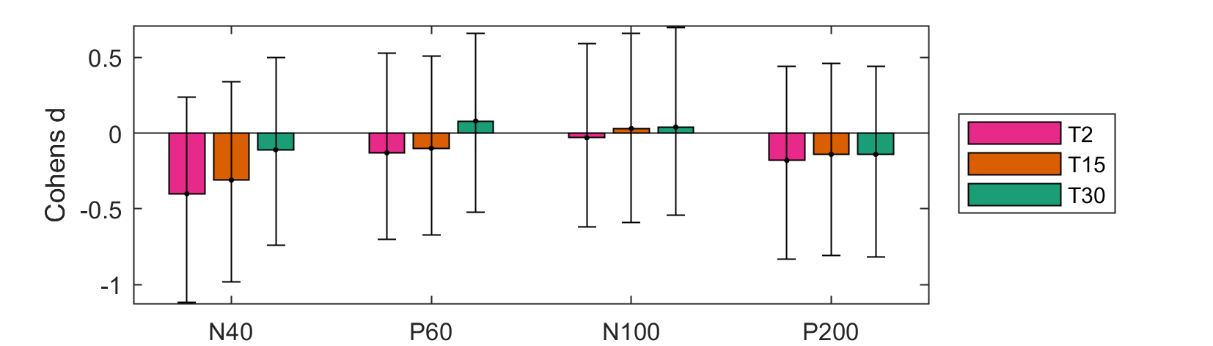

S10. Global scalp analyses across time

When assessing pre-post differences within the sham condition, one significant positive cluster (p = 0.014, 199-240 ms) was identified*.* Over time, there were significant differences in amplitude between pre-sham and T2, T15 and T30 (p=0.003, p=0.021 and p=0.009, respectively), with pre-sham amplitude more positive than at all post-sham time points. Scalp topography revealed that these differences were more pronounced in central and centroparietal sensors (Supplementary Material, Fig. S10A).

Within the cTBS condition, permutation tests showed a significant positive cluster (p = 0.008, 117-143 ms), with T15 and T30 TEP amplitudes more positive (a decrease in the negative values) than pre-cTBS. Topographical representation in this latency range showed that these differences in amplitudes were more pronounced in fronto-central and central sensors (Supplementary Material, Fig. S10B). Using cluster-based permutation analysis, we did not find any pre-post differences in TEP amplitudes for iTBS (all p > 0.025, two-tailed test).

A) Sham

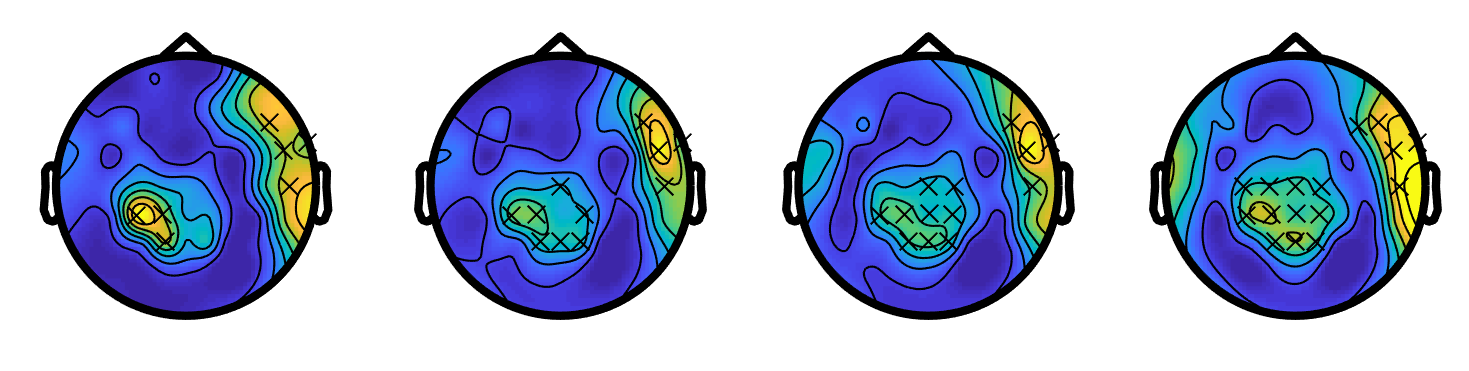

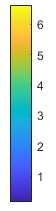

F-value

200ms	210ms	220ms	230ms

B) cTBS

F-value

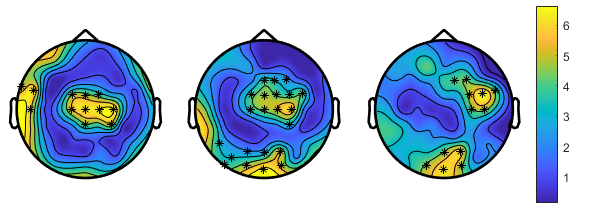

120ms	130ms	140ms

**Fig S10.** Significant changes in TEP amplitude at global level. Post more negative (yellow) or positive (blue) than pre. (‘*’ indicate significant clusters with p<0.01, ‘X’ p<0.025)
